# Using local convolutional neural networks for genomic prediction

**DOI:** 10.1101/2020.05.12.090118

**Authors:** Torsten Pook, Jan Freudenthal, Arthur Korte, Henner Simianer

## Abstract

The prediction of breeding values and phenotypes is of central importance for both livestock and crop breeding. With increasing computational power and more and more data to potentially utilize, Machine Learning and especially Deep Learning have risen in popularity over the last few years. In this study, we are proposing the use of local convolutional neural networks for genomic prediction, as a region specific filter corresponds much better with our prior genetic knowledge of traits than traditional convolutional neural networks. Model performances are evaluated on a simulated maize data panel (n = 10,000) and real Arabidopsis data (n = 2,039) for a variety of traits with the local convolutional neural network outperforming both multi layer perceptrons and convolutional neural networks for basically all considered traits. Linear models like the genomic best linear unbiased prediction that are often used for genomic prediction are outperformed by up to 24%. Highest gains in predictive ability was obtained in cases of medium trait complexity with high heritability and large training populations. However, for small dataset with 100 or 250 individuals for the training of the models, the local convolutional neural network is performing slightly worse than the linear models. Nonetheless, this is still 15% better than a traditional convolutional neural network, indicating a better performance and robustness of our proposed model architecture for small training populations. In addition to the baseline model, various other architectures with different windows size and stride in the local convolutional layer, as well as different number of nodes in subsequent fully connected layers are compared against each other. Finally, the usefulness of Deep Learning and in particular local convolutional neural networks in practice is critically discussed, in regard to multi dimensional inputs and outputs, computing times and other potential hazards.

## 1 INTRODUCTION

The prediction of breeding values and phenotypes is of central importance for both livestock and crop breeding. Obtaining accurate estimates of breeding values at an earlier time point can impact the decision on which individuals and lines to keep in a breeding programs, reducing the generation cycle and therefore leading to higher genomic gains per year (Schaeffer, 2006). Optimizing breeding schemes is of key importance for overcoming the global challenges of feeding a planet with a rising Human population (Foley et al., 2011).

The most commonly applied method for the prediction of breeding values and phenotypes consider a mixed model or bayesian linear models (Meuwissen et al., 2001; Gianola et al., 2009; Erbe et al., 2012). With the availability of genomic data, traditional methods that rely on parental relationships and pedigrees have been replaced by genomic evaluations in which the pedigree-based relationship matrix has been replaced by a variant construced from genomic data (VanRaden, 2008). Currently, variations of this approach have been successfully implemented in both animal (Hayes et al., 2009; Hayes and Goddard, 2010; Gianola and Rosa, 2015) and plant breeding (Jannink et al., 2010; Albrecht et al., 2011; Nakaya and Isobe, 2012; Heslot et al., 2015). As breeding values are additive by design, most of these models only account for additive single marker effects, but adaptations to account for dominance and epistatic interactions have been proposed (Da et al., 2014; Jiang and Reif, 2015; Martini, 2017) and are regularly applied for the prediction of phenotypes. In recent years the use of deep learning (DL) and in particular artifical neural networks (ANN) have become more and more populuar in a variety of fields in genetics (Eraslan et al., 2019). This is further enhanced by a variety of available open-source libraries like Keras (Chollet, 2015) and Tensorflow (Abadi et al., 2016) which combine options for a simple and flexible set up of ANNs with a highly efficient computational back end.

The transition from traditional mixed models and bayesian linear models to the use of DL for genomic prediction seems like a natural next step, as reflected by a variety of recent studies (Bellot et al., 2018; Waldmann, 2018; Ma et al., 2018; Montesinos-López et al., 2019; Pérez-Enciso and Zingaretti, 2019; Azodi et al., 2019; Khaki and Wang, 2019) reporting peformance of multi-layer perceptrons (MLP) and convolutional neural networks (CNN) for a variety of traits in both humans and a wide set of livestock and crop species. The common result in those studies is that traditionally applied statistical methods such as genomic best linear unbiased prediction (GBLUP) or methods from the bayesian alphabet (Meuwissen et al., 2001; Gianola et al., 2009; Erbe et al., 2012) lead to similar or slightly higher predictive ability. In cases for which improvements were achieved, either very specific trait architectures are considered (Waldmann, 2018), improvements are not consistent across traits (Bellot et al., 2018; Montesinos-López et al., 2019) or additional data like environmental information is used (Khaki and Wang, 2019). For most traits considered in those studies, the best performing ANNs are usually MLPs with one or sometimes two fully-connected layers (FCL) between input and output layer (Bellot et al., 2018; Montesinos-López et al., 2019). Predictive ability obtained with CNNs is usually similar or even slightly worse (Bellot et al., 2018) with best performing models using very small filters. On first glance, this might be surprising since in other fields one of the biggest reasons for the rise of ANNs is attributed to the use of CNNs and convolutional layers (CL) (Krizhevsky et al., 2012; Goodfellow et al., 2016; Ubbens and Stavness, 2017). One must consider here that SNP arrays only contain markers and no full genome sequence. Therefore, a specific sequence of alleles on a SNP-chip in one region can not be linked to the same sequence of alleles in another region. As traditional CLs are directly assuming this, naive use of a CL does not make much sense from a modelling perspective. Therefore, we here propose the use of local convolutional layers (LCL) to allow for the use of region specific filters while still maintaining the positive features of a CL like the massively reduced number of parameters in the model. Region specific filter means that in contrast to a CL, parameters of the layers can vary based on the region, e.g. for a toy example given in Figure 1 *a, d, g* can be different whereas a CNN architecture would result in *a* = *d* = *g*. In the following, the performance of local convolutional neural networks (LCNN) is compared to both traditional methods for genomic prediction and other more commonly applied ANN architectures.

**Figure 1.**
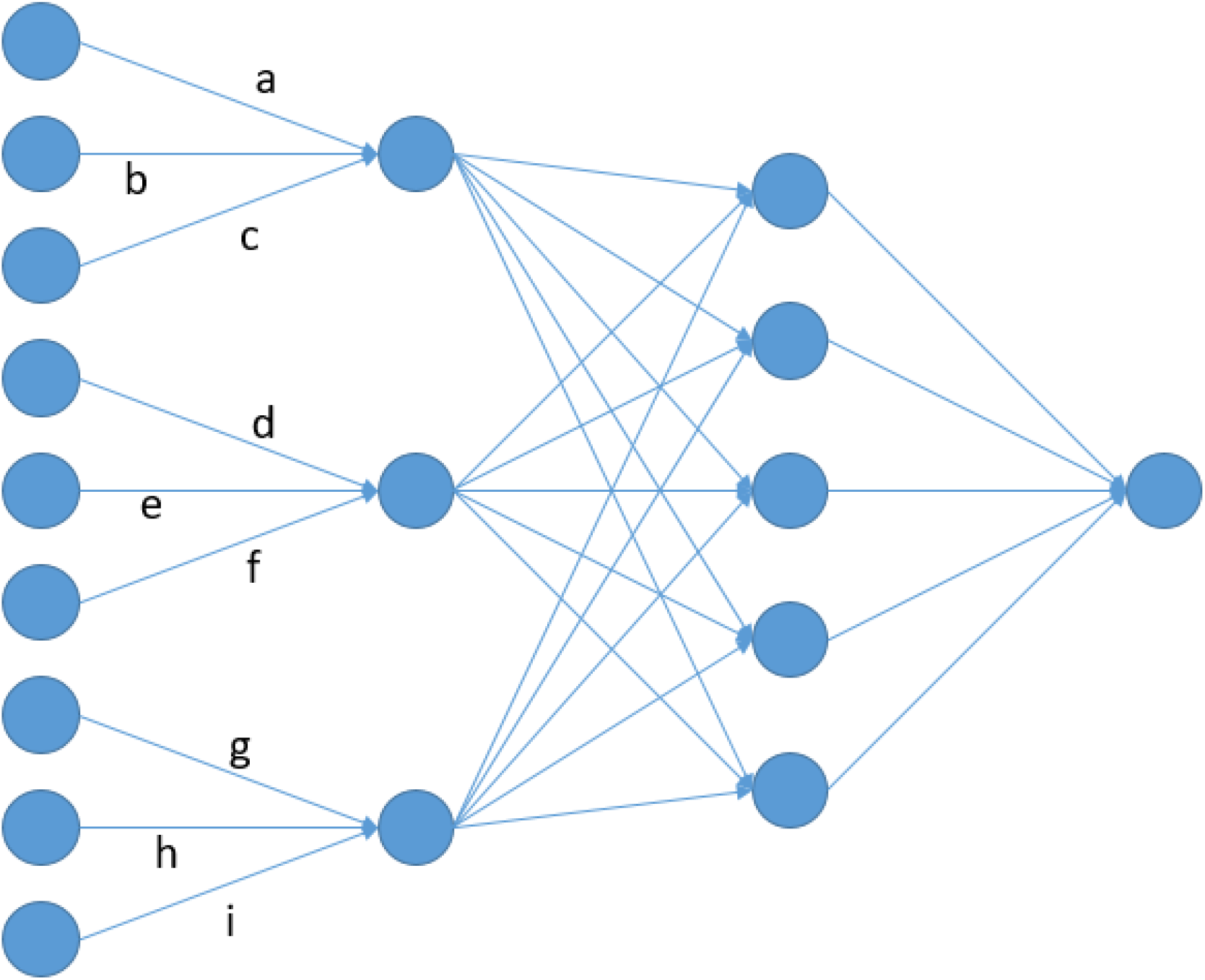
Node architecture of an LCNN containing a LCL with window size and stride of 3 and a FCL with 5 nodes.

## 2 MATERIAL AND METHODS

### 2.1 Material

As a first dataset, a simulated data panel containing 10,000 maize lines that were genotyped at 34,595 SNPs with 17 traits of different trait complexities ranging from traits with 10 additive single locus QTL to traits caused by epistatic interaction between potentially physically linked QTL was considered. Individual effect sizes were drawn from a gaussian, gamma and binomial distribution. The dataset was generated based on simulations in the R-package MoBPS (Pook et al., 2020) and original genotypes stem from 501 doubled haploid lines of the European maize landrace Kemater Landmais Gelb that were genotyped via the Affymetrix Axiom Maize Genotyping Array (Unterseer et al., 2014) and reduced via LD pruning in PLINK (Purcell et al., 2007). The interested reader is referred to Hölker et al. (2019) for details on the data generation procedure. The R-code used to generate the 10,000 individuals and the 17 traits in MoBPS is available in Supplementary File S1. For each trait, residuals variances were varied to obtain traits with a heritability *h*^2^ of 0.1, 0.5, 0.8 and 1.

As a second dataset, a real data panel from the 1001 genomes project of Arabidopsis thaliana (Alonso-Blanco et al., 2016) was considered. After quality control, filtering for minor allele frequency and LD pruning, we reduced the available 10.7 M SNPs to 180k SNPs for 2,029 lines. Tests were conducted for 50 different traits that were available and contained measurements for between 83 and 468 lines (Atwell et al., 2010; Li et al., 2010; Meijón et al., 2014; Strauch et al., 2015; Seren et al., 2016). The interested reader is referred to Freudenthal (2020) for details on the data preparation steps.

Scripts used to perform the model fitting in Keras (Chollet, 2015) are available in Supplementary File S2 and S3. The R-packages rrBLUP (Endelman, 2011) and BGLR (Pérez and de los Campos, 2014) were used for fitting of the linear models.

### 2.2 Design of the neural network

For all tested ANNs, the SNP dataset with genotypes coded as 0,1,2 was used as the input layer and (centered) phenotypes were used as the output layer. In genomic prediction and in particular when using an ANN, the number of parameters is substantially higher than the number of individuals that can be used for the model fitting. Thus, leading to n ≪ p problems (Fan et al., 2014). In this study, we will compare four main classes of models:

1. Linear models (LM)
2. Multi-layer perceptrons (MLP)
3. Convolutional neural networks (CNN)
4. Local convolutional neural networks (LCNN)

For the class of LMs a variety of models have been proposed. The most frequently applied linear model in todays’ applications is the genomic best linear unbiased predictor (GBLUP, (Meuwissen et al., 2001)) that is using a mixed model in which the variance of the random effect is given by a relationship matrix like the one propsed by VanRaden (2008). An alternative to this are methods typically referred to as the bayesian alphabet (Gianola et al., 2009; de los Campos et al., 2013) that perform bayesian linear regression with prior assumptions on the individual marker variance, e.g. BayesA is using a scaled-t-distribution as the prior. In particular for phenotype prediction, there are a variety of other genomic relationship matrices for the mixed model have been proposed to account for non-additive effects. The extended genomic best linear unbiased predictor (EGBLUP, (Martini, 2017)) is designed to assign linear effects to specific marker combination and therefore is able to include epistatic interactions into the mixed model.

All three other classes describe different types of ANNs. Here, we define the class of MLP as ANNs that only contain FCLs. In CNN / LCNN we are using an additional single CL / LCL in front of the FCLs without any use of pooling. For all three ANN classes we tested different layer designs ranging from just one up to three FCLs with varying number of nodes. For the CNN and LCNN we also tested different filters for the convolutional layer ranging from windows size and strides between 3 and 40 with potential overlap between windows. For all models the relu function was used as the activation function with an adam optimizer (Kingma and Ba, 2014) to minimize the mean squared errors with a dropout rate of 0.3 after each layer (Chollet, 2015; Goodfellow et al., 2016). Changes to activation function, optimizer, dropout rate and target function were also tested but only had neglectable effects and are therefore are neglected in the following.

Models are compared based on their predictive ability on the test set (80% of the samples used for model fitting, 20% as a test set), and we define the predictive ability as the correlation of the predicted genomic values and their phenotypes.

### 2.3 Size and structure of the training data

A well-known problem of ANNs is that overfitting can occur after a high number of training epochs (Goodfellow et al., 2016). Therefore, we split the 8,000 samples used for model fitting for the simulated maize data into 7,000 samples used for the actual training of the model (training set) and 1,000 samples that are just used to determine at what state training should be stopped (validation set). After each epoch the predictive ability of the model was derived based on the validation set and the best performing model from up to 50 epochs was used as the final model. In the same way the validation set can also be used to derive the ideal architecture of the ANN.

To further investigate the impact of the size of the training population, we considered different sizes of the training data (100, 250, 500, 1,000, 2,000, 3,000, 4,000, 6,000, 8,000). The size of the validation set was adapted based on the size of the training data (20, 50, 100, 200, 300, 400, 500, 750, 1,000), as with smaller data panels an higher impact of the validation set was observed. For the Arabidopsis data, the data used for model fitting was split into 80% used for model fitting and 20% used for validation. As the training data for most of the Arabidopsis traits was already extremely small, a second study was conducted in which a fixed number of 25 epochs was performed with no validation set and therefore larger training set.

All tests for the simulated data / Arabidopsis data were repeated 25 / 100 times respectively, with randomly sampled training and test sets.

## 3 RESULTS

### 3.1 Comparison between model types

In the following, we will report results for a representative model from each of the three ANN class:

1. MPL: 2 FCL with 64 nodes
2. CNN: CL with kernel size and stride 10 + 2 FCL with 64 nodes
3. LCNN: LCL with kernel size and stride 10 + 2 FCL with 64 nodes

Minor improvements were obtained by tweaking parameter settings for selected traits but overall tendencies of predictive ability across filter size and number of nodes as well as layers were stable. More details on differences will be provided for the LCNN at the end of the results section. For the LMs there was no clear best model for all traits. We will consider GBLUP as the baseline, but also report results for BayesA (Meuwissen et al., 2001) and the EGBLUP model (Martini, 2017). As results for effect sizes drawn from gaussian, gamma and binomial distribution were very similar, we will only report results for the effect sizes drawn from a gaussian distribution.

### 3.2 Simulated data

In the following, we will first report results for the traits with a simulated heritability of 0.5. In the purely additive setting with just 10 underlying QTL the highest predictive ability was obtained with the LCNN (0.666), outperforming the other three baseline models by around 0.03-0.04 (Table 1, Figure 2 **(A)**). When increasing the number of QTL to 1,000, differences between LCNN (0.606) and the other three baseline models increased to around 0.06-0.09 (Table 1, Figure 2 **(B)**). The BayesA model led to similar preditive ability (0.660) as the LCNN for 10 QTL but was outperformed (0.538) in case of 1,000 underlying QTL. Even though the simulated traits had a purely additive genetic background, the EGBLUP model led to very similar or even slightly higher predictive ability as the GBLUP model. A potential reason for this could be “phantome epistatis” (de los Campos et al., 2019) as the use of pair-wise marker interactions could lead to a better overall representation of haplotype similarities.

**Figure 2.**
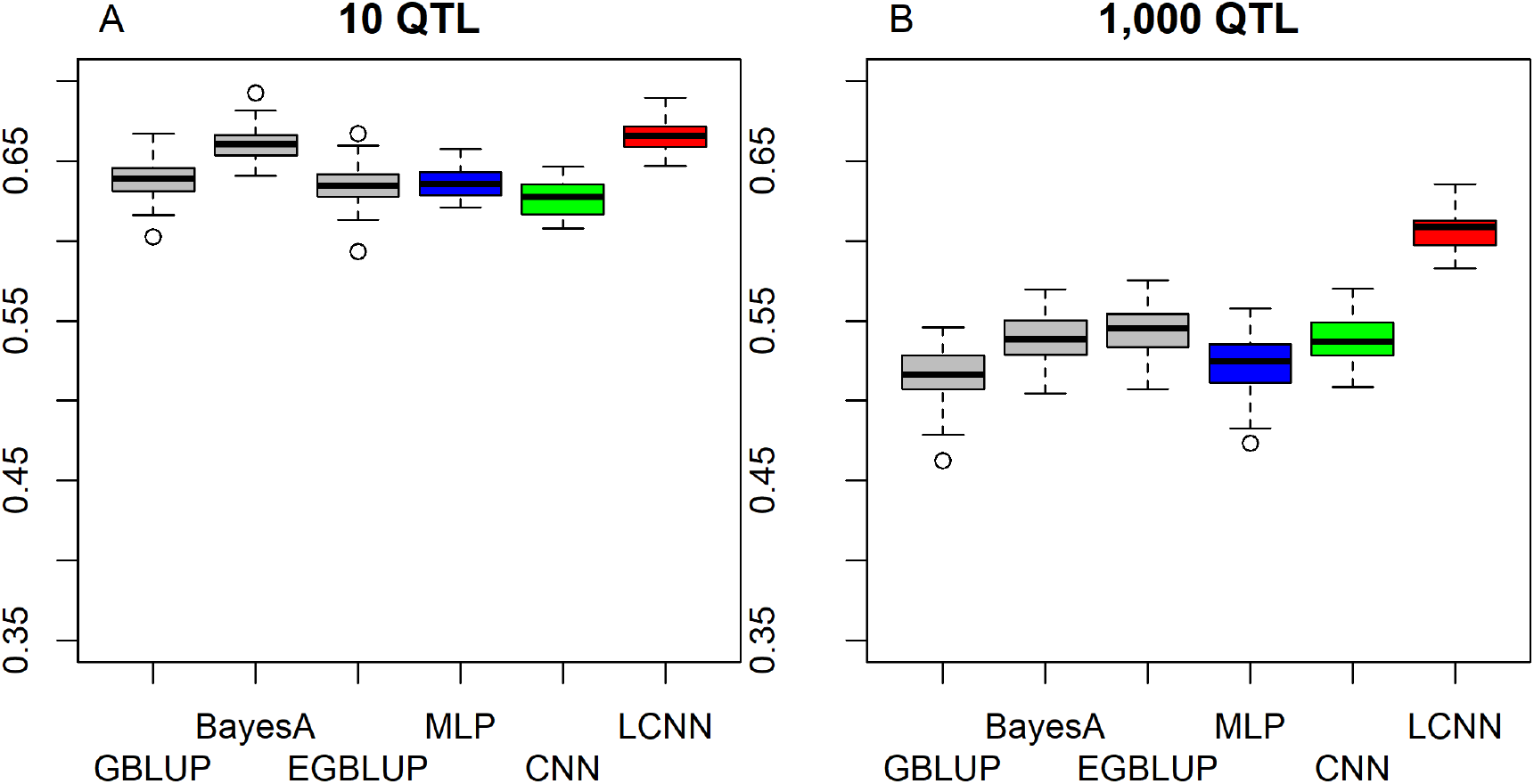
Predictive ability of different methods for genomic prediction for a simulated trait with 10 (A) and 1,000 (B) purely additive QTL.

**Table 1.** Predictive ability for the different models on different traits with *h*^2^ = 0.5.

| Trait architecture | GBLUP | BayesA | EGBLUP | MPL | CNN | LCNN |
| --- | --- | --- | --- | --- | --- | --- |
| 10 additive QTL | 0.639 | 0.660 | 0.635 | 0.637 | 0.627 | 0.666 |
| 1,000 additive QTL | 0.516 | 0.538 | 0.543 | 0.524 | 0.538 | 0.606 |
| 10 epistatic QTL | 0.511 | 0.527 | 0.519 | 0.503 | 0.491 | 0.572 |
| 1,000 epistatic QTL | 0.416 | 0.414 | 0.448 | 0.395 | 0.403 | 0.401 |
| 10 locally linked epistatic QTL | 0.488 | 0.501 | 0.529 | 0.504 | 0.544 | 0.625 |
| 1,000 locally linked epistatic QTL | 0.524 | 0.523 | 0.541 | 0.519 | 0.517 | 0.510 |

When considering a purely epistatic trait architecture with 10 underlying QTL, differences between the LCNN and the other three baseline models are also around 0.06-0.08 (Figure 3 **(A)**), whereas results in the case of 1,000 underlying QTL were very similar for all four baseline models (Table 1, Figure 3 **(B)**) with the GBLUP model (0.416) leading to slightly higher predictive ability (0.01-0.02). In case the underlying QTL of the epistatic trait were played on physically linked markers to imitate a trait caused by local interactions in a gene, both the LCNN and CNN obtained higher predictive ability when only 10 QTL were involved in the trait, whereas the MLP and GBLUP performed worse (Figure 4 **(A)**). The relative differences between LCNN (0.625) and GBLUP (0.488) were here highest among all considered cases. In the case of 1000 locally linked underlying QTL, results of the four baseline models were again very similar with GBLUP performing about 0.01 better than the ANNs (Figure 4 **(B)**). In all cases of epistatic QTL, the use of the EGBLUP model led to higher predictive abilities than GBLUP. For both cases of 10 underlying epistatic QTL the LCNN model was still superior, whereas the EGBLUP model was best for traits with 1,000 underlying epistatic QTL.

**Figure 3.**
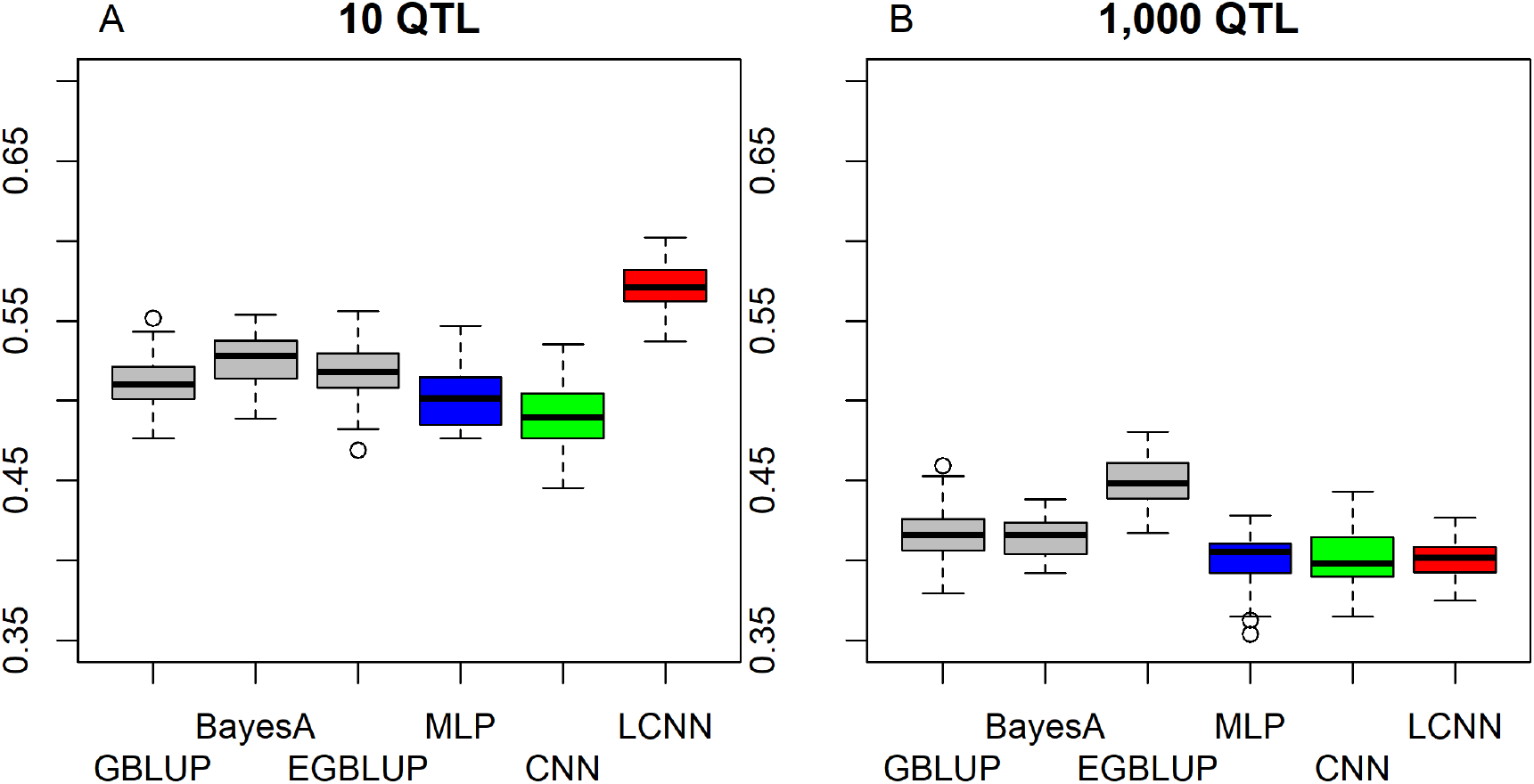
Predictive ability of different methods for genomic prediction for a simulated trait with 10 (A) and 1,000 (B) purely non-linked epistatic QTL.

**Figure 4.**
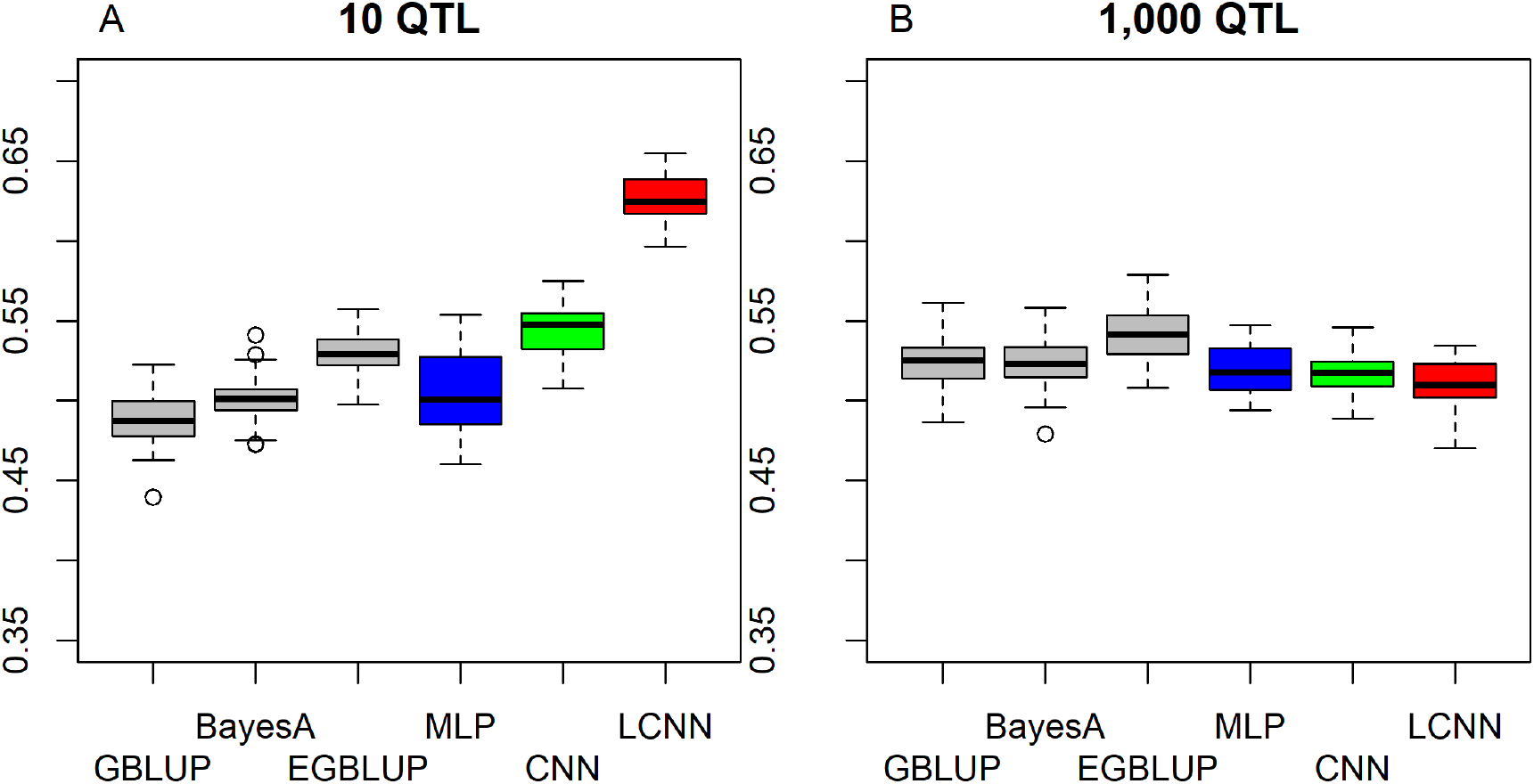
Predictive ability of different methods for genomic prediction for a simulated trait with 10 (A) and 1,000 (B) purely non-linked epistatic QTL.

When considering traits with varying heritability, higher overall predictive ability for traits with higher heritability was observed. This was even the case after standardizing the predictive ability by dividing with the squared root of the heritablity as this is the highest achievable correlation between phenotypes and estimated breeding values (Figure 5). Overall obtained standardized predictive ability for the additive traits are higher and close to the maximum in the case of 10 additive underlying QTL (Figure 5 **(A)**). In particular for cases of high heritablity, the LCNN is outperforming all other models for both the additive trait with 1,000 QTL and the epistatic traits with 10 QTL (Figure 5 **(B,C,E)**). For the epistatic traits with 1,000 QTL all models are on a similar level for all considered heritablities (Figure 5 **(D,F)**).

**Figure 5.**
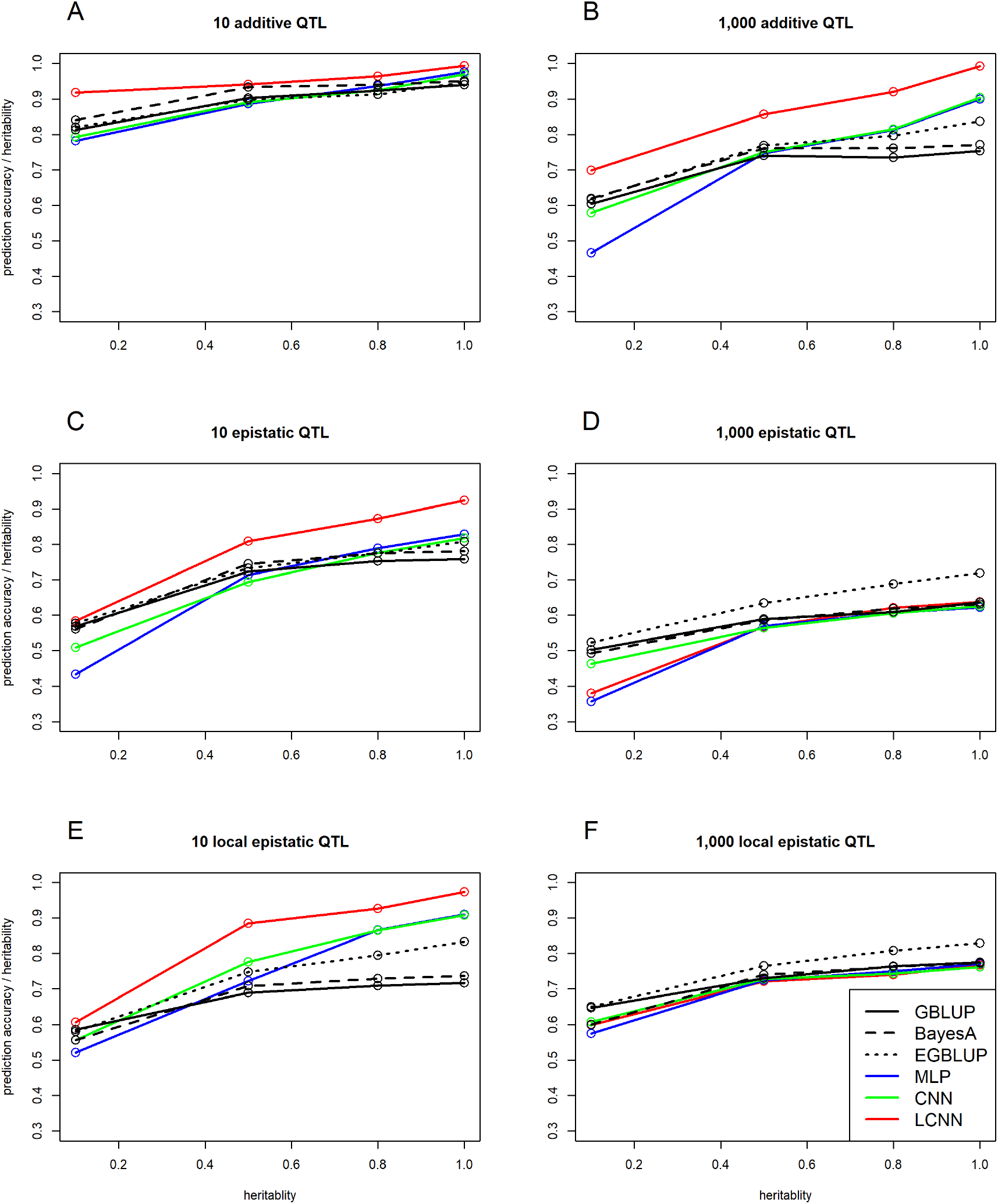
Predictive ability of the LCNN compared to the GBLUP model in relation to the trait heritability for the purely additive (A/B), epistatic (C/D) and physically linked epistatic (E/F) trait with 10/1,000 underlying QTL.

When comparing the predictive ability depending on the number of individuals used for training, we observed worse performance of all three classes of ANN models relative to GBLUP for small training sets. In particular training sets of size 100 and 250 led to massive drops in predictive ability. Of the three ANN classes considered, the LCNN performed best and with the exception of the epistatic traits with 1,000 underlying QTL was at least close to the performance of GBLUP. In particular for traits with 1,000 purely additive QTL and 10 epistatic QTL the increase in predictive ability was substantially higher than in all three considered linear models (Figure 6). As ANNs are known to be extremely data hungry (Goodfellow et al., 2016) this should not be that surprising. Overall, the ANN architectures with less layers and parameters were less affected by the reduced size of the training set.

**Figure 6.**
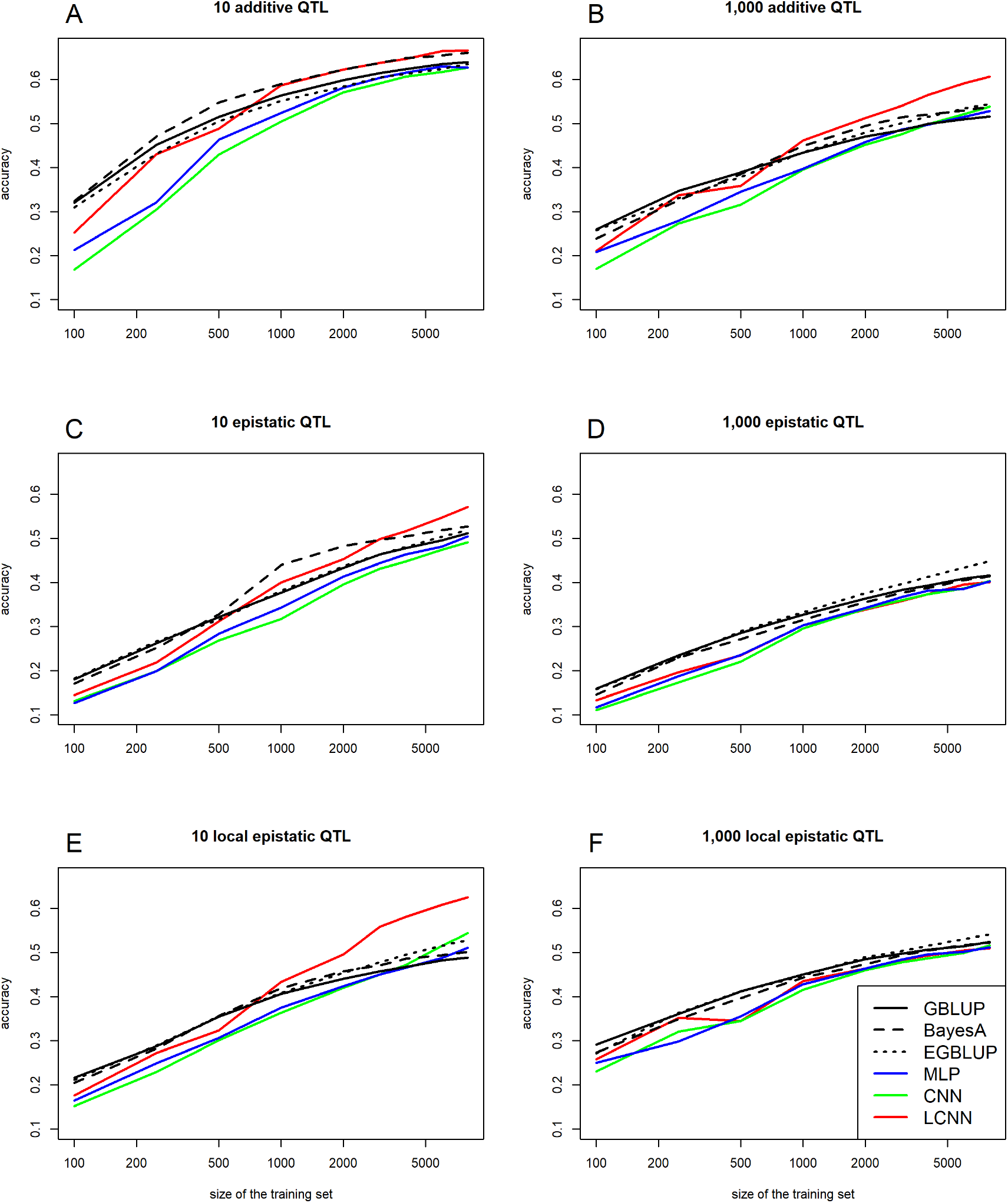
Predictive ability of the representative LCNN model and BayesA depending on the size of the training set for purely additive (A/B), epistatic (C/D) and physically linked epistatic (E/F) trait with 10/1,000 underlying QTL.

### 3.3 Comparison between LCNN models

When comparing different layer designs for the LCNN, we observed small, but still significant differences between the different model architectures. In particular for purely additive traits, larger window sizes (WS) in the LCL led to higher accuracies (WS 5: 0.603; WS 10: 0.606; WS 20: 0.616), whereas the stride had neglectable impact (Figure 7). In regard to the design of the following FCLs, we observed increased predictive abilities when using a high number nodes (128 / 256) per layer (Figure 8). Differences between the highest obtain predictive ability for the different number of layers were neglectable, as long as at least one FCL was used.

**Figure 7.**
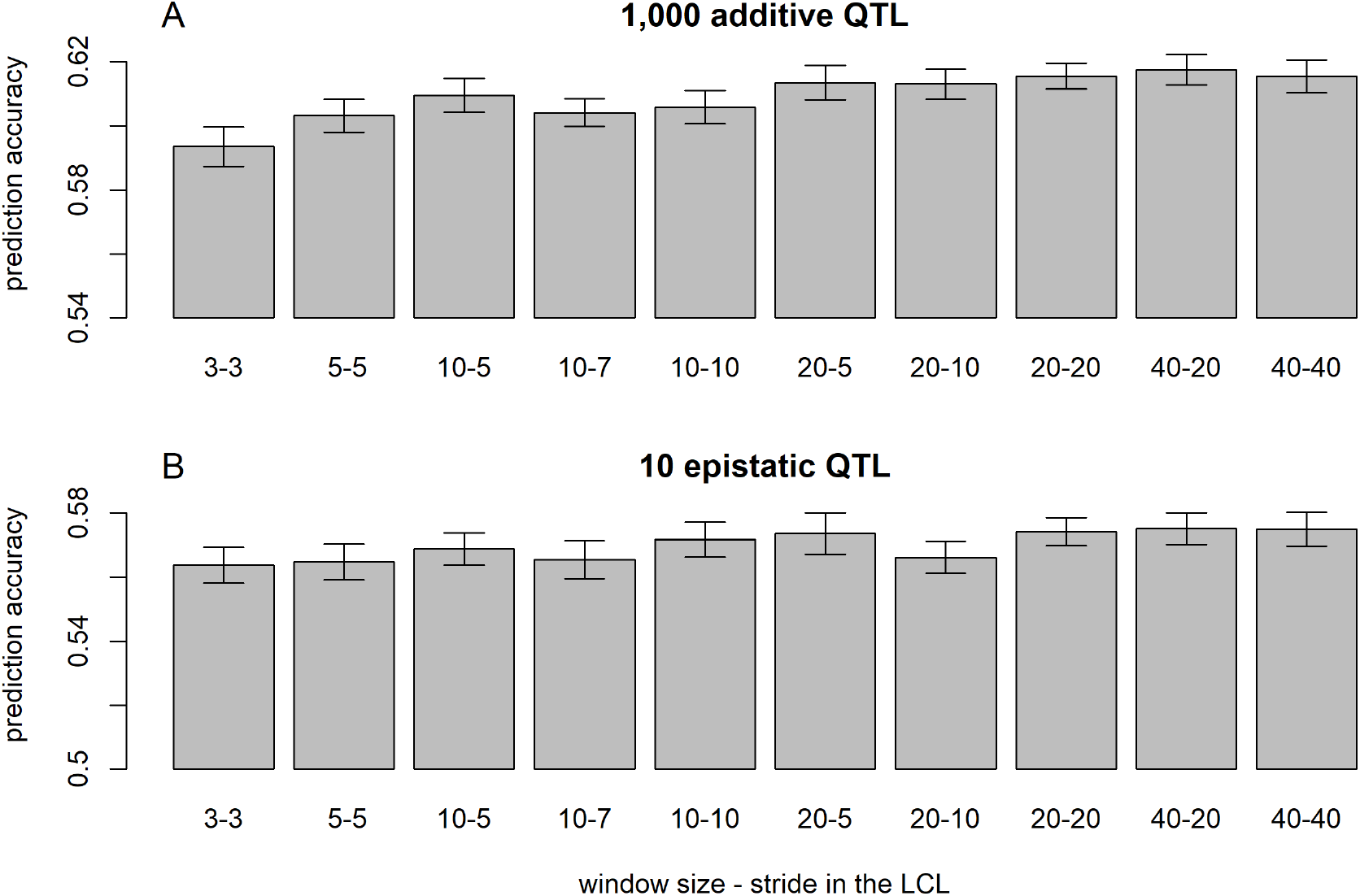
Predictive ability of different layer designs of the LCNN with modifications to the LCL for the purely additive trait with 1,000 QTL (A) and the epistatic trait with 10 QTL (B).

**Figure 8.**
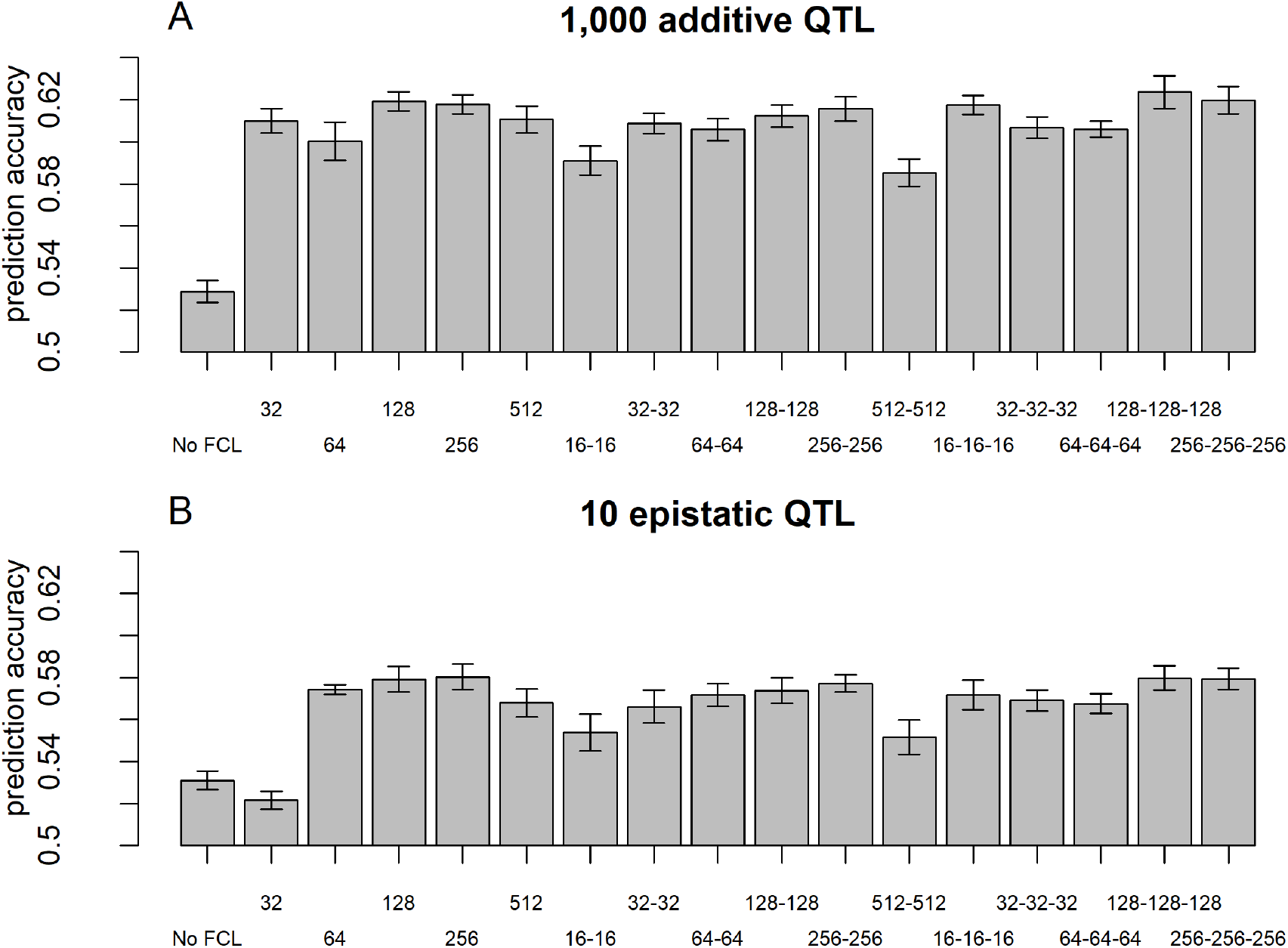
Predictive ability of different layer designs of the LCNN with modifications to the FCLs for the purely additive trait with 1,000 QTL (A) and the epistatic trait with 10 QTL (B).

### 3.4 Arabidopsis data

When comparing the different ANN models for the Arabidopsis dataset, the highest average predictive ability was observed for the LCNN model (0.340) compared to 0.316 for the MLP and 0.312 for the CNN (Table 2). All three ANNs were however outperformed by the three linear models (GBLUP, BayesA, EGBLUP). The differences between the ANNs and the linear models is decreasing for traits with higher number of individuals used in the training set. Whereas differences for traits with less than 100 individuals on average were 0.078 between GBLUP and the LCNN, this differences is reduced to 0.037 / 0.021 for traits with more than 100 / 250 lines in the training set (Table 2). The variance in obtained predictive ability was highest for MLP (0.031) and CNN (0.031) compared to the LCNN (0.029) and lowest for the linear models (0.024). Note that no traits with more than 468 phenotyped lines were considered here and gains in the simulated data were typically only obtained for training set with at least 1,000 lines (Figure 6). When not using a validation set the overall accuracies are going up for all three considered ANN architectures and performances are more similar to GBLUP (Table 3, Figure 9). One exception to this is the trait FT field which resulted in extremely unstable models for all three ANNs with 20% of all trained models leading to basically zero predictive ability and on average 55% lower predictive ability. Details on the predictive ability of the individual traits and the number of phenotypes considered for each trait are given in Supplementary S4. Additional minor improvement were obtained by modifying the layer design for the FCLs after the LCNN. The interested reader is referred to Freudenthal (2020) for details on those extended benchmarking tests. Note that after trait-specific model architecture tunings in Freudenthal (2020) higher predictive ability with the LCNN compared to GBLUP were obtained for 33 of the 52 traits with *h*^2^ > 0.5 were obtained, whereas only 27 of the 93 traits with *h*^2^ < 0.5 benefited from the use of an LCNN compared to GBLUP.

**Figure 9.**
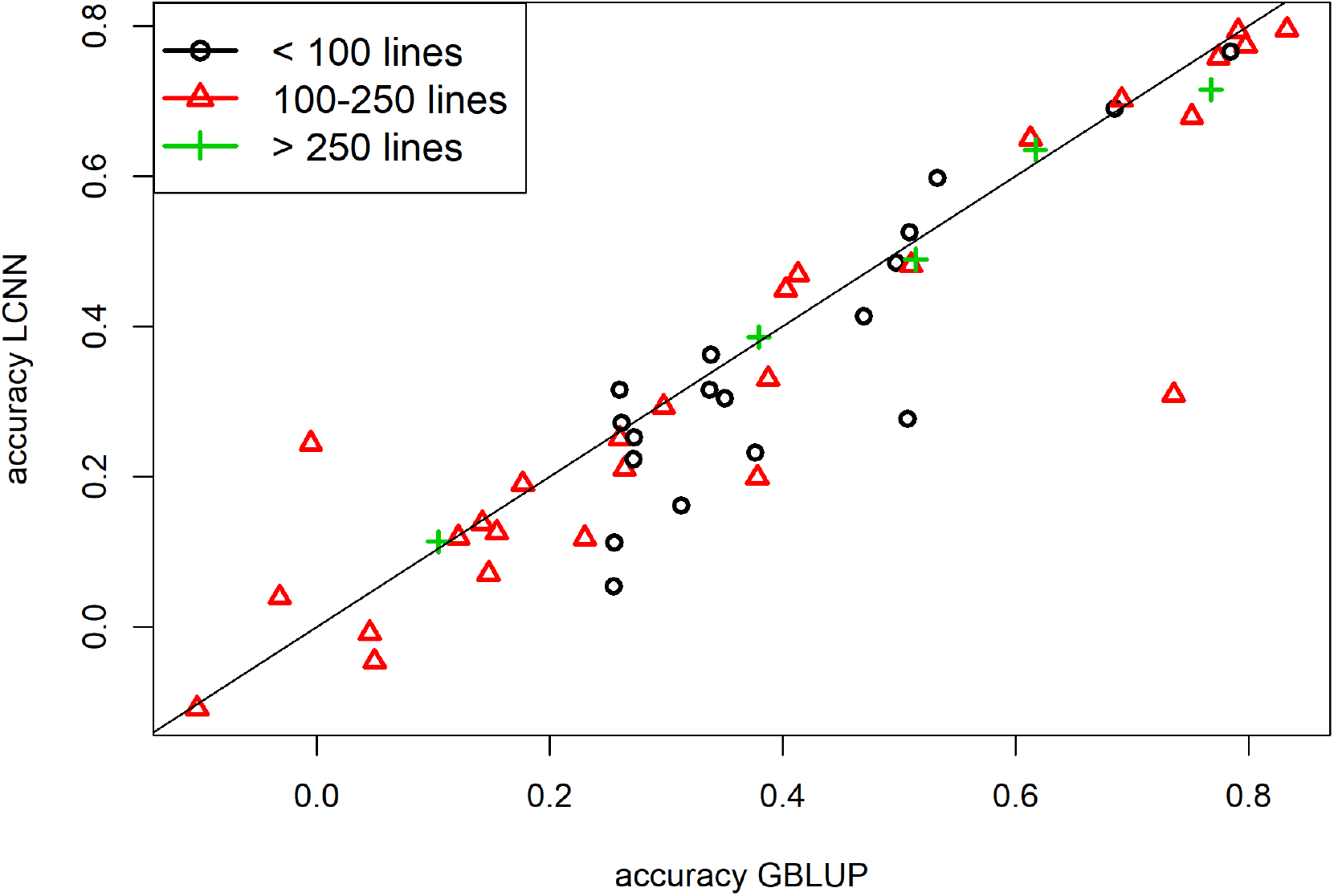
Predictive ability for GBLUP and the LCNN model for the different arabidopsis traits in relation to the size of the training set and no validation set.

**Table 2.** Average predictive ability for the different models for the Arabidopsis traits in relation to the size of the training set.

| Trait architecture | GBLUP | BayesA | EGBLUP | MPL | CNN | LCNN |
| --- | --- | --- | --- | --- | --- | --- |
| Average predictive ability (all) | 0.390 | 0.382 | 0.382 | 0.316 | 0.312 | 0.340 |
| Average predictive ability (training set < 100) | 0.404 | 0.390 | 0.399 | 0.300 | 0.299 | 0.326 |
| Average predictive ability (100 < training set < 250) | 0.364 | 0.358 | 0.354 | 0.318 | 0.311 | 0.327 |
| Average predictive ability (training set > 250) | 0.477 | 0.477 | 0.472 | 0.358 | 0.370 | 0.456 |

**Table 3.**
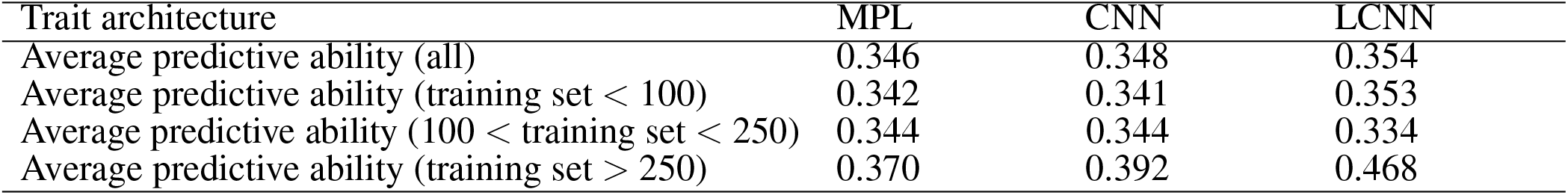
Average predictive ability for the different models for the Arabidopsis traits in relation to the size of the training set and no validation set.

| Trait architecture | MPL | CNN | LCNN |
| --- | --- | --- | --- |
| Average predictive ability (all) | 0.346 | 0.348 | 0.354 |
| Average predictive ability (training set < 100) | 0.342 | 0.341 | 0.353 |
| Average predictive ability (100 < training set < 250) | 0.344 | 0.344 | 0.334 |
| Average predictive ability (training set > 250) | 0.370 | 0.392 | 0.468 |

## 4 DISCUSSION

A common misconception of ANNs is that they are handled and used as black-boxes, leading to back propagation of causal variants and fundamental model design questions to be second order problems. Note that the baseline MLP models used in our tests results in a model with 2.2 million parameters and 8,000 individuals and thereby leading to potential massive problems of overparametrization (Fan et al., 2014). The use of a CL is reducing this problems substantially with our baseline CNN “only” needing 225,610 parameters. However, CLs assign effects to specific sequences of input variants. As the same sequence of markers on an array in different segments of the genome is usually not linked in any way, this modelling approach does not really make sense from a genetics perspective and thus makes it potentially more difficult to obtain a good model fit. The LCNN fixes these issues by introducing region-specific filters. This increases the number of required parameters in the model slightly (260,195), but still is a massive improvement in terms of number of parameters compared to the MLP. When working with whole genome sequence the use of CNNs has shown to be quite useful (Washburn et al., 2019). However, whole-genome sequence data does not reflect the currently available standard for genomic prediction, as no significant gains in most applications are reported when using more than just low to medium density SNP arrays (Ober et al., 2012; Erbe et al., 2013), generating such sequence data is still costly (Schwarze et al., 2018) and problems of even higher overparametrization can arise here.

As shown by the results above, the use of a LCNN can massively improve the accuracy of genomic prediction compared to frequently applied ANNs architectures like MLPs and CNNs for both simulated and real datasets and in particular for traits with small training sets. In the case of the simulated data, improvements compared to linear models like GBLUP were obtained for both simulated purely additive and purely epistatic traits. However, for the real Arabidopsis data panel with at most a couple of hundred lines per phenotype, average predictive ability was slightly reduced as in particular for traits with small training sets, predicitive abilities was substantially lower for the ANNs when a validation set was used. However when using a set number of training epoch and no validation set were almost on the level of GBLUP. Note however that the use of no validation set, requires prior knowledge on a reasonable number of training epochs and model architecture, therefore leading to potential model instability. The use of a LCNN was an improvement compared to more commonly applied ANN architectures (MLPs/CNNs) in both cases. The variance in predictive ability for the ANN models was slightly higher than for the linear models, but the differences were not large enough to cause major concerns in regard to model stability of the ANNs.

Whereas significantly higher numbers of genotyped lines in the setting of plant breeding are not realistic, even larger populations with potentially millions of animals are available in livestock breeding. As in particular for traits of medium complexity (1,000 additive QTL & 10 epistitatic QTL) substantially gains for the LCNN compared to all other models were obtained, these results indicate high potentially for genomic prediction in such traits as traditional linear models tend to reach a plateau in predictive ability (Erbe et al., 2013). However, a potential problem for the use in animal breeding is that for all considered individuals the same inputs have to be provided and therefore requiring the genotyping of all individuals. Particularly to be mentioned here is that there is no direct equivalent to single step GBLUP (Legarra et al., 2009; Christensen and Lund, 2010) to combine pedigree and genotype data in a joint relationship matrix up till now. Furthermore, one needs to consider that breeding values are additive by design and even if higher predictive ability is obtained with non-additive models, this will not necessarily result in higher genetic gains under a random mating environment (Martini et al., 2017). This leads us to conclude that ANNs (and in fact epistatic models like EGBLUP in general) are much better suited for the prediction of phenotypes than breeding values (Martini et al., 2017).

A further potential application for the use of ANNs that is in particular relevant for plant breeding is the inclusion of other omics, environmental data or even information about weather condictions, as ANNs are very flexible in their design and it is relatively easy to add additional input and/or output layers to an existing model. Computing times and model complexity in the framework of ANNs are far less affected by such additional inputs than GBLUP-based models (Gillberg et al., 2019). As such ANNs typically contain separate layers for each input dimension and those are concarnated in later steps, the use of an LCL for the SNP-based inputs should be highly beneficial for such applications.

When deciding between the use of ANNs and traditional linear models there are however more things to consider than just the plain predictive abilities. This particularly includes potential economic issues, as the use of ANNs would at this moment require genotyping of all individuals, and required conceptional changes to modern breeding programs as terms like reliability do not have a direct equivalent in ANNs and therefore among others require changes in the design of selection indicies (Hazel and Lush, 1942; Miesenberger, 1997). Additional work in checking if higher predictive ability also translate into higher genomic gains is a further topic that needs to be investigated, as even the use of epistatic models have shown to not always lead to higher gain, despite higher predictive ability (Martini et al., 2017).

Nonetheless, we can conclude that there is considerable potential in the use of ANNs and in particular LCNN in genomic prediction when working with large individual numbers and high heritability. and/or additional input dimensions like other omics. We would expect the highest potential of ANNs to be especially relevant with more complex input and output layers, as present when considering different omics (Li et al., 2019), weather data (Gillberg et al., 2019) or prediction across environments (Freudenthal, 2020) as inputs, or multiple correlated traits as outputs (Lyra et al., 2017). Accounting for such input/outputs in the traditional models, even in a linear way, was shown to be extremely costly from a computational side and oftentimes does not significantly improve results (Calus and Veerkamp, 2011). With generation of such datasets becoming cheaper and widely available, we would expect the use of DL techniques to be of increasing importances for quantitative genetics and in particular genomic prediction in the near future.

## AUTHOR CONTRIBUTIONS

TP lead the development of the methodology, performed the analysis and wrote the initial manuscript. JF, AK, HS provided critical feedback to both analysis and the manuscript. HS supervised the study. All authors read and approach the final manuscript.

## ACKNOWLEDGMENTS

We acknowledge support by the Open Access Publication Funds of the Göttingen University. The authors further thank the German Federal Ministry of Education and Research (BMBF) for the funding of our project (MAZE – “Accessing the genomic and functional diversity of maize to improve quantitative traits”; Funding ID: 031B0195).

## DATA AVAILABILITY STATEMENT

Publicly available datasets were analyzed in this study. This data can be found here:

https://arapheno.1001genomes.org/

https://link.springer.com/article/10.1007/s00122-019-03428-8.

