## Supplementary material for "Using local convolutional neural networks for genomic prediction": File S1: File_S1.R

```
### Generation of simulated data panels

## Import real genotype data from Hoelker et al 2019
data <- as.matrix(read.table("MAZE_data_50k_SNP.txt"))

## Effect-structure
set.seed(42)

add <- list()
mult <- list()
# 10 additive QTL - Same Size
sign <- rbinom(10,1,0.5)
add[[1]] <- cbind(sample(1:nrow(data), 10), 1,sign*2-1, 0, -sign*2-1)
mult[[1]] <- NULL

# 1000 additive QTL - Same Size
sign <- rbinom(1000,1,0.5)
add[[2]] <- cbind(sample(1:nrow(data), 10), 1,sign*2-1, 0, -sign*2-1)
mult[[2]] <- NULL

# 10 two-marker QTL
add[[3]] <- NULL
mult[[3]] <- cbind(sample(1:nrow(data), 10), 1, sample(1:nrow(data), 10), 1,matrix(rgamma(10*9,1), ncol=9) )

# 1000 two-marker QTL
add[[4]] <- NULL
mult[[4]] <- cbind(sample(1:nrow(data), 1000), 1, sample(1:nrow(data), 1000), 1,matrix(rgamma(1000*9,1), ncol=9) )

# 10 two-marker QTL (highly linked)
add[[5]] <- NULL
dist <- sample(1:5,10, replace=TRUE)
pos1 <- sample(1:(nrow(data)-max(dist)), 10)
mult[[5]] <- cbind( pos1, 1,pos1+ dist, 1,matrix(rgamma(10*9,1), ncol=9) )

# 1000 two-marker QTL (highly linked)
add[[6]] <- NULL
dist <- sample(1:5,1000, replace=TRUE)
pos1 <- sample(1:(nrow(data)-max(dist)), 1000)
mult[[6]] <- cbind( pos1, 1,pos1+ dist, 1,matrix(rgamma(1000*9,1), ncol=9) )


# 10 two-marker QTL (linked)
# close 1-25
add[[7]] <- NULL
dist <- sample(1:25,10, replace=TRUE)
pos1 <- sample(1:(nrow(data)-max(dist)), 10)
mult[[7]] <- cbind(pos1, 1,pos1+ dist,1, matrix(rgamma(10*9,1), ncol=9) )

# 1000 two-marker QTL (linked)
# close 1-25
add[[8]] <- NULL
dist <- sample(1:25,1000, replace=TRUE)
pos1 <- sample(1:(nrow(data)-max(dist)), 1000)
mult[[8]] <- cbind( pos1, 1,pos1+ dist,1, matrix(rgamma(1000*9,1), ncol=9) )


# 10 two-marker QTL
add[[9]] <- NULL
mult[[9]] <- cbind( sample(1:nrow(data), 10), 1, sample(1:nrow(data), 10), 1,matrix(rnorm(10*9), ncol=9) )

# 1000 two-marker QTL
add[[10]] <- NULL
mult[[10]] <- cbind(sample(1:nrow(data), 1000), 1, sample(1:nrow(data), 1000), 1,matrix(rnorm(1000*9), ncol=9) )

# 10 two-marker QTL (highly linked)
add[[11]] <- NULL
dist <- sample(1:5,10, replace=TRUE)
pos1 <- sample(1:(nrow(data)-max(dist)), 10)
mult[[11]] <- cbind( pos1, 1,pos1+ dist, 1,matrix(rnorm(10*9), ncol=9) )

# 1000 two-marker QTL (highly linked)
add[[12]] <- NULL
dist <- sample(1:5,1000, replace=TRUE)
pos1 <- sample(1:(nrow(data)-max(dist)), 1000)
mult[[12]] <- cbind( pos1, 1,pos1+ dist, 1,matrix(rnorm(1000*9), ncol=9) )


# 10 two-marker QTL (linked)
add[[13]] <- NULL
dist <- sample(1:25,10, replace=TRUE)
pos1 <- sample(1:(nrow(data)-max(dist)), 10)
mult[[13]] <- cbind( pos1, 1,pos1+ dist, 1,matrix(rnorm(10*9), ncol=9) )

# 1000 two-marker QTL (linked)
add[[14]] <- NULL
dist <- sample(1:25,1000, replace=TRUE)
pos1 <- sample(1:(nrow(data)-max(dist)), 1000)
mult[[14]] <- cbind( pos1, 1,pos1+ dist, 1,matrix(rnorm(1000*9), ncol=9) )

# 1000 two-marker QTL
add[[15]] <- NULL
mult[[15]] <- cbind( sample(1:nrow(data), 1000), 1, sample(1:nrow(data), 1000), 1,matrix(rbinom(1000*9,1,1/9), ncol=9) )

# 1000 two-marker QTL (highly linked)
add[[16]] <- NULL
dist <- sample(1:5,1000, replace=TRUE)
pos1 <- sample(1:(nrow(data)-max(dist)), 1000)
mult[[16]] <- cbind( pos1, 1,pos1+ dist, 1,matrix(rbinom(1000*9,1,1/9), ncol=9) )

# 1000 two-marker QTL (linked)
add[[17]] <- NULL
dist <- sample(1:25,1000, replace=TRUE)
pos1 <- sample(1:(nrow(data)-max(dist)), 1000)
mult[[17]] <- cbind( pos1, 1,pos1+ dist, 1,matrix(rbinom(1000*9,1,1/9), ncol=9) )

# For simplicity assume just a single chromosome of length 10 Morgan
library(MoBPS)
pop1 <- creating.diploid(dataset = data/2, chromosome.length = 10,
                        real.bv.add = add, real.bv.mult = mult, miraculix=TRUE)

pop2 <- breeding.diploid(pop1, selection.size=pop1$info$size[1,], breeding.size = c(10000,0))


bv <- get.bv(pop2, gen=2)
geno <- get.geno(pop2, gen=2)

bv_var <- diag(var(t(bv)))
bv_mean <- rowMeans(bv)
bv <- (bv-bv_mean) / sqrt(bv_var)

pheno01 <- bv + rnorm(length(bv), 0, sd=3)
pheno05 <- bv + rnorm(length(bv), 0, 1)
pheno08 <- bv + rnorm(length(bv), 0, sd=0.5)

write.table(file="Pheno08.txt", t(pheno08), quote=FALSE, col.names = FALSE, row.names = FALSE)
write.table(file="Pheno05.txt", t(pheno05), quote=FALSE, col.names = FALSE, row.names = FALSE)
write.table(file="Pheno01.txt", t(pheno01), quote=FALSE, col.names = FALSE, row.names = FALSE)
write.table(file="Pheno10.txt", t(bv), quote=FALSE, col.names = FALSE, row.names = FALSE)
write.table(file="geno_small.txt", geno[,1:1000], quote=FALSE, col.names = FALSE, row.names = FALSE)
write.table(file="geno.txt", geno, quote=FALSE, col.names = FALSE, row.names = FALSE)
```
