## Supplementary material for "Using local convolutional neural networks for genomic prediction": File S2

May 11, 2020

In [ ]:
```
import numpy as np
import pandas as pd
from keras import backend as K
import keras
import random
import tensorflow as tf
import multiprocessing as multi
from keras.models import Sequential
from keras.layers import Dense, Dropout, Activation, LocallyConnected1D, Embedding
from keras.layers import GaussianNoise, Flatten, LSTM, Convolution1D
from keras.optimizers import SGD
from keras import regularizers
from scipy.stats.stats import pearsonr
from keras.models import load_model
import matplotlib.pyplot as plt

dataframe2= pd.read_csv('Genetic_Datasets/Maize_sim/Pheno05.txt',sep=" ",header=None)
dataframe1= pd.read_csv('Genetic_Datasets/Maize_sim/geno.txt',sep=" ",header=None)
epoch = 50
batch = 32
dataset2= dataframe2.values
dataset1= dataframe1.values
X2 = dataset1
X = X2.transpose()
X = np.expand_dims(X, axis=2)

for index in range(0,25):
    for x in range(0,17):

        Y = dataset2[:,x]
        settingnr = 1

        model = Sequential()
        model.add(LocallyConnected1D(1,10, strides=10, input_shape=(34595,1)))
        model.add(Flatten())
        Dropout(0.3)
        model.add(Dense(64, kernel_initializer='normal', activation='relu'))
        Dropout(0.3)
```

```
model.add(Dense(64, kernel_initializer='normal', activation='relu'))
Dropout(0.3)
model.add(Dense(1, kernel_initializer='normal'))
model.compile(loss='mse', optimizer='adam')

training = random.sample(range(0,10000), 8000)
test = np.delete(range(0,10000), training, 0)
train_vali = random.sample(training, 1000)
del1 = []
for i in range(0,1000):
    del1.append(int(train_vali[i]))
for i in range(0,2000):
    del1.append(int(test[i]))
train_train = np.delete(range(0,10000),del1,0)
X_train = X[train_train,]
Y_train = Y[train_train]
X_vali = X[train_vali,]
Y_vali = Y[train_vali]
X_test = X[test,]
Y_test = Y[test]
best = float('inf')
train_error = []
vali_error = []
test_error = []

for i in range(epoch):
    model.fit(X_train, Y_train, epochs=1, batch_size=batch)
    train_cur = model.evaluate(X_train,Y_train)
    vali_cur = model.evaluate(X_vali,Y_vali)
    test_cur = model.evaluate(X_test,Y_test)
    train_error.append(float(train_cur))
    vali_error.append(float(vali_cur))
    test_error.append(float(test_cur))
    if best>vali_cur:
        best = vali_cur
        model.model.save('my_model'+str(settingnr)+'.h5', overwrite=True)
        best_epoch = i
model2 = load_model('my_model'+str(settingnr)+'.h5')

Y_hat = model2.predict(X)
Y3 = []
for i in train_train:
    Y3.append(float(Y[i]))
Y_hat3 = []
for i in train_train:
    Y_hat3.append(float(Y_hat[i]))
Y4 = []
for i in train_vali:
```

```
    Y4.append(float(Y[i]))
Y_hat4 = []
for i in train_vali:
    Y_hat4.append(float(Y_hat[i]))
Y5 = []
for i in test:
    Y5.append(float(Y[i]))
Y_hat5 = []
for i in test:
    Y_hat5.append(float(Y_hat[i]))

training_cor = pearsonr(Y_hat3, Y3)
vali_cor = pearsonr(Y_hat4, Y4)
test_cor = pearsonr(Y_hat5, Y5)

string1 = 'train_cor'+str(settingnr)+'.txt'
string2 = 'vali_cor'+str(settingnr)+'.txt'
string3 = 'test_cor'+str(settingnr)+'.txt'
string4 = 'train_error'+str(settingnr)+'.txt'
string5 = 'vali_error'+str(settingnr)+'.txt'
string6 = 'test_error'+str(settingnr)+'.txt'
string7 = 'best'+str(settingnr)+'.txt'
file = open(string1,"a")
file.write('%f' % training_cor[0] + ' ')
file.close()
file = open(string2,"a")
file.write('%f' % vali_cor[0] + ' ')
file.close()
file = open(string3,"a")
file.write('%f' % test_cor[0] + ' ')
file.close()
file = open(string4,"a")
for i in range(epoch):
    file.write('%f' % train_error[i] + ' ')
file.close()
file = open(string5,"a")
for i in range(epoch):
    file.write('%f' % vali_error[i] + ' ')
file.close()
file = open(string6,"a")
for i in range(epoch):
    file.write('%f' % test_error[i] + ' ')
file.close()
file = open(string7,"a")
file.write('%f' % best_epoch + ' ')
file.close()
K.clear_session()
gc.collect()
```
