## Supplementary material for "Using local convolutional neural networks for genomic prediction": File S3

May 11, 2020

```
In [ ]: import numpy as np
        import pandas as pd
        from keras import backend as K
        import keras
        import random
        import tensorflow as tf
        import multiprocessing as multi
        from keras.models import Sequential
        from keras.layers import Dense, Dropout, Activation, LocallyConnected1D, Embedding
        from keras.layers import GaussianNoise, Flatten, LSTM, Convolution1D
        from keras.optimizers import SGD
        from keras import regularizers
        from scipy.stats.stats import pearsonr
        from keras.models import load_model
        import matplotlib.pyplot as plt

        for settingnr in range(0,51):

            dataframe2=pd.read_csv('AB_data/Pheno'+str(settingnr)+'.txt',sep=" ",header=None)
            dataframe1=pd.read_csv('AB_data/Geno'+str(settingnr)+'.txt',sep=" ",header=None)
            epoch = 25
            batch = 32
            dataset2= dataframe2.values
            dataset1= dataframe1.values
            X2 = dataset1
            X = X2.transpose()
            X = np.expand_dims(X, axis=2)

            for index in range(0,100):

                Y = dataset2[:,2]
                nt = X.shape[0]
                nt1 = int(0.8 * nt)
                nt2 = int(0.2 * nt)
                nv = int(nt1 * 0.2)
                ns = X.shape[1]

                model = Sequential()
```

```
model.add(LocallyConnected1D(1,10, strides=10, input_shape=(ns,1)))
model.add(Flatten())
Dropout(0.3)
model.add(Dense(64, kernel_initializer='normal', activation='relu'))
Dropout(0.3)
model.add(Dense(64, kernel_initializer='normal', activation='relu'))
Dropout(0.3)
model.add(Dense(1, kernel_initializer='normal'))
model.compile(loss='mse', optimizer='adam')

training = random.sample(range(0,nt), nt1)
test = np.delete(range(0,nt), training, 0)
train_vali = random.sample(training, nv)
del1 = []
for i in range(0,nt2):
    del1.append(int(test[i]))
train_train = np.delete(range(0,nt),del1,0)
X_train = X[train_train,]
Y_train = Y[train_train]
X_vali = X[train_vali,]
Y_vali = Y[train_vali]
X_test = X[test,]
Y_test = Y[test]
best = float('inf')
train_error = []
vali_error = []
test_error = []
for i in range(epoch):
    model.fit(X_train, Y_train, epochs=1, batch_size=batch)
    train_cur = model.evaluate(X_train,Y_train)
    vali_cur = model.evaluate(X_vali,Y_vali)
    test_cur = model.evaluate(X_test,Y_test)
    train_error.append(float(train_cur))
    vali_error.append(float(vali_cur))
    test_error.append(float(test_cur))
    if i==(epoch-1):
        best = vali_cur
        model.model.save('Ara_my_model'+str(settingnr)+'.h5', overwrite=True)
        best_epoch = i
model2 = load_model('Ara_my_model'+str(settingnr)+'.h5')

Y_hat = model2.predict(X)
Y3 = []
for i in train_train:
    Y3.append(float(Y[i]))
Y_hat3 = []
for i in train_train:
    Y_hat3.append(float(Y_hat[i]))
```

```
Y4 = []
for i in train_vali:
    Y4.append(float(Y[i]))
Y_hat4 = []
for i in train_vali:
    Y_hat4.append(float(Y_hat[i]))
Y5 = []
for i in test:
    Y5.append(float(Y[i]))
Y_hat5 = []
for i in test:
    Y_hat5.append(float(Y_hat[i]))

training_cor = pearsonr(Y_hat3, Y3)
vali_cor = pearsonr(Y_hat4, Y4)
test_cor = pearsonr(Y_hat5, Y5)

string1 = 'AraLCNN_train_cor'+str(settingnr)+'.txt'
string2 = 'AraLCNN_vali_cor'+str(settingnr)+'.txt'
string3 = 'AraLCNN_test_cor'+str(settingnr)+'.txt'
string4 = 'AraLCNN_train_error'+str(settingnr)+'.txt'
string5 = 'AraLCNN_vali_error'+str(settingnr)+'.txt'
string6 = 'AraLCNN_test_error'+str(settingnr)+'.txt'
string7 = 'AraLCNN_best'+str(settingnr)+'.txt'
file = open(string1,"a")
file.write('%f' % training_cor[0] + ' ')
file.close()
file = open(string2,"a")
file.write('%f' % vali_cor[0] + ' ')
file.close()
file = open(string3,"a")
file.write('%f' % test_cor[0] + ' ')
file.close()
file = open(string4,"a")
for i in range(epoch):
    file.write('%f' % train_error[i] + ' ')
file.close()
file = open(string5,"a")
for i in range(epoch):
    file.write('%f' % vali_error[i] + ' ')
file.close()
file = open(string6,"a")
for i in range(epoch):
    file.write('%f' % test_error[i] + ' ')
file.close()
file = open(string7,"a")
file.write('%f' % best_epoch + ' ')
file.close()
```

```
K.clear_session()
gc.collect()
```
